# Deep-Learning-Based Accelerated and Noise-Suppressed Estimation (DANSE) of quantitative Gradient Recalled Echo (qGRE) MRI metrics associated with Human Brain Neuronal Structure and Hemodynamic Properties

**DOI:** 10.1101/2021.09.10.459810

**Authors:** Sayan Kahali, Satya V.V.N. Kothapalli, Xiaojian Xu, Ulugbek S. Kamilov, Dmitriy A. Yablonskiy

## Abstract

**Purpose:** To introduce a Deep-Learning-Based Accelerated and Noise-Suppressed Estimation (DANSE) method for reconstructing quantitative maps of biological tissue cellular-specific, *R*2*t*^*^ and hemodynamic-specific, *R*2′ from Gradient-Recalled-Echo (GRE) MRI data with multiple gradient-recalled echoes.

**Methods:** DANSE method adapts supervised learning paradigm to train a convolutional neural network for robust estimation of *R*2*t*^*^ and *R*2′ maps free from the adverse effects of macroscopic (*B*_0_) magnetic field inhomogeneities directly from the GRE magnitude images without utilizing phase images. The corresponding ground-truth maps were generated by means of a voxel-by-voxel fitting of a previously-developed biophysical quantitative GRE (qGRE) model accounting for tissue, hemodynamic and *B*_0_-inhomogeneities contributions to GRE signal with multiple gradient echoes using nonlinear least square (NLLS) algorithm.

**Results:** We show that the DANSE model efficiently estimates the aforementioned brain maps and preserves all features of NLLS approach with significant improvements including *noise-suppression* and *computation speed* (from many hours to seconds). The noise-suppression feature of DANSE is especially prominent for data with SNR characteristic for typical GRE data (SNR~50), where DANSE-generated *R*2*t*^*^ and *R*2′ maps had three times smaller errors than that of NLLS method.

**Conclusions:** DANSE method enables fast reconstruction of *magnetic-field-inhomogeneity-free* and *noise-suppressed* quantitative qGRE brain maps. DANSE method does not require any information about field inhomogeneities during application. It exploits spatial patterns in the qGRE MRI data and previously-gained knowledge from the biophysical model, thus producing clean brain maps even in the environments with high noise levels. These features along with fast computational speed can lead to broad qGRE clinical and research applications.

## Introduction

Magnetic resonance imaging (MRI) has become a powerful non-invasive tool to investigate the human brain structure in health and disease. Numerous MRI pulse sequences are used in clinical practice to create so-called “weighted” images (e.g. diffusion-weighted, T1- or T2-weighted) tuned to emphasize different brain structures and pathologies. More detail information can be potentially obtained from quantitative MRI approaches (e.g. measuring water molecule diffusion, T1 or T2 relaxation time parameters) that are more sensitive to different aspects of brain tissue microstructure than weighted approaches. While weighted MRI usually allows fast reconstruction of images from collected k-space data, the quantitative MRI approaches more often than not require long reconstruction times using sophisticated computer programs and are also more sensitive to noise in the data than the weighted MRI. These issues often create barriers for implementing quantitative MRI for clinical applications.

In this paper, we are considering one of these techniques, a quantitative Gradient Recalled Echo (qGRE) MRI (Ulrich & Yablonskiy, 2016). qGRE MRI (Ulrich & Yablonskiy, 2016), is based on the gradient recalled echo sequence with multiple gradient recalled echoes, biophysical model of GRE signal decay (Yablonskiy, 1998; Yablonskiy & Haacke, 1994) and algorithms for data analysis allowing separating contributions of the tissue specific (*R*2*t*^*^), blood-oxygen-level-dependent (*R*2′) and adverse effects of macroscopic magnetic field inhomogeneities (D. A. Yablonskiy, A. L. Sukstanskii, J. Luo, & X. Wang, 2013a) from the total R2*=1/T2* relaxation rate parameter of the GRE signal. Importantly, the quantitative relationships established in (Wen, Goyal, Astafiev, Raichle, & Yablonskiy, 2018) between brain cellular composition and the *R*2*t*^*^ metric of the quantitative Gradient Recalled Echo (qGRE) MRI (Ulrich & Yablonskiy, 2016) shows that the *R*2*t*^*^ subcomponent of the R2* is associated with the neuronal properties of human brain:

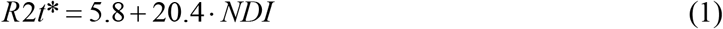

where *R*2*t*^*^ is measured in inverse seconds and NDI is a neuronal density index (dimensionless parameter ranging from 0 for tissue void of neurons to 1 for tissue that would consist only from neuronal cells). In addition, the *R*2′ subcomponent of R2* can be used to measure concentration of deoxyhemoglobin per unit tissue volume by means of the relationship (Ulrich & Yablonskiy, 2016):

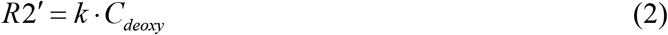

where parameter *k* is defined below in the Methods section, Eq. (8).

The qGRE technique has already proven useful in studying of healthy brain aging (Zhao, Wen, Cross, & Yablonskiy, 2016), loss of neurons at early stages of Alzheimer disease (Kothapalli et al., 2021; Zhao et al., 2017), multiple sclerosis (Xiang, Wen, Cross, & Yablonskiy, 2019) and traumatic brain injury (Astafiev et al., 2019). While qGRE sequence can be implemented on any commercial MRI scanner and requires less than 10 minutes to acquire high resolution data, the data analysis currently cannot be done on the scanner as it requires hours of computing time to generate *R*2*t*^*^ and *R*2′ maps using fitting routines based on non-linear least squares (NLLS) approaches.

Advent of introducing machine learning approaches to the field of medical image reconstruction (Aggarwal, Mani, & Jacob, 2019; Castiglioni et al., 2021; Eo et al., 2018; Hammernik et al., 2018; Knoll et al., 2019; Knoll et al., 2020; Schlemper, Caballero, Hajnal, Price, & Rueckert, 2018; Sriram et al., 2020; Sun, Li, & Xu, 2016; Yaman et al., 2020; Yang et al., 2018; Zhu, Liu, Cauley, Rosen, & Rosen, 2018) opened new opportunities for generating quantitative MRI data instead of using NLLS-based fitting routines. The methods developed so far mostly focused on training Artificial Neural Network (ANN) for a voxel-by-voxel parameter estimations (Domsch et al., 2018; Hubertus et al., 2019). In particular, (Domsch et al., 2018) demonstrated improved accuracy in estimating oxygen extraction fraction (OEF) from GRE data and theoretical model (Yablonskiy, 1998) based on ANN-fitting.

Recently we have demonstrated (Torop et al., 2020) that using deep learning approach allows reconstruction of combined R2* maps in a matter of seconds with improved image quality and reduced noise effects. However, separate generating quantitative maps of R2t* and *R*2′ is a more challenging problem as it requires data with significantly higher signal-to-noise ratio (SNR) (X. Wang, Sukstanskii, & Yablonskiy, 2013; Zhao et al., 2016) not always available from experimental data. From this perspective, using deep neural network instead of ANN provides clear advantage in dealing with noisy data (Torop et al., 2020) and also reducing computation time. Moreover, we have developed a deep learning framework (Xu et al., 2021) which performs motion correction on complex qGRE images and enables reconstruction of high-quality motion-free quantitative R2* maps.

This paper presents a fully supervised Deep-Learning-Based Accelerated and Noise-Suppressed Estimation (DANSE) method for generating qGRE metrics associated with human brain neuronal structure and hemodynamic properties. Our method is based on Convolutional Neural Network (CNN) which is trained on qGRE MR images reflecting signal decay. The network architecture is similar to our recently published RoAR method (Torop et al., 2020). However, the ground truth maps were generated by fitting the biophysical model (Ulrich & Yablonskiy, 2016) to experimental GRE data with multiple gradient echoes using nonlinear least square algorithm. Moreover, the biophysical model accounts for the macroscopic magnetic field inhomogeneities (background field gradients), thus producing *R*2*t*^*^ and *R*2′ maps essentially free from the background field gradients –induced artifacts. Importantly, the DANSE method generates the *R*2*t*^*^ and *R*2′ brain maps directly from the qGRE MR magnitude images without the need for utilizing k-space data and/or phase images.

Our results show that applying deep leaning DANSE approach to qGRE MRI data allows generating *R*2*t*^*^ and *R*2′ maps with significant acceleration of reconstruction times (from many hours to seconds) with simultaneous improvement in image quality (by reducing contribution of noise to reconstructed data).

## Methods

In this study, we used previously published 26 qGRE brain imaging data collected from 26 healthy volunteers (age range 26-76) using a Siemens 3T Trio MRI scanner and a 32-channel phased-array head coil (Zhao et al., 2016). All studies were approved by local IRB of Washington University. All volunteers provided informed consent.

### Biophysical model of qGRE MRI signal formation

To identify brain cellular structure and hemodynamic properties we use the quantitative Gradient Recalled Echo (qGRE) MRI method (Ulrich & Yablonskiy, 2016). The qGRE method is based on 3D gradient recalled echo MRI sequence with multiple gradient echoes and theoretical model that describes GRE signal relaxation properties. In the framework of this approach, the GRE signal dependence on the echo time *TE* is presented in the following equation ((Yablonskiy, 1998), (Ulrich & Yablonskiy, 2016))

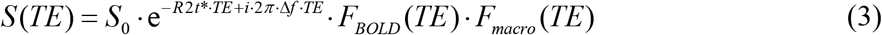

where *TE* is the gradient echo time, *S*_*0*_ is the signal amplitude, R2t* is the tissue-specific (t- stands for tissue) transverse relaxation rate constant (describing GRE signal decay in the absence of BOLD effect), Δ*f* is the frequency shift (dependent on tissue structure and also macroscopic magnetic field created mostly by tissue/air interfaces), function *F*_*BOLD*_*(TE)* describes GRE signal decay due to the presence of blood vessel network with deoxygenated blood (veins and the part of capillaries adjacent to veins) (Yablonskiy & Haacke, 1994), and function *F*_*macro*_*(TE)* accounts for the adverse effects of macroscopic magnetic field inhomogeneities. Herein, *F(TE)* is calculated by means of a voxel spread function (VSF) method (D. A. Yablonskiy, A. L. Sukstanskii, J. Luo, & X. Q. Wang, 2013b). In this paper for the BOLD model (Yablonskiy & Haacke, 1994) we use the following expression (Ulrich & Yablonskiy, 2016)

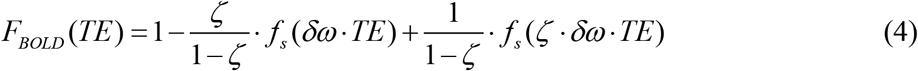

that more completely accounts for the presence of large veins. In Eq. (4), *ζ* is the deoxygenated cerebral blood volume (dCBV) fraction (deoxyhemoglobin-containing part of the total blood volume, i.e. veins and adjacent to them portion of the capillary bed), and the characteristic frequency *δω* is:

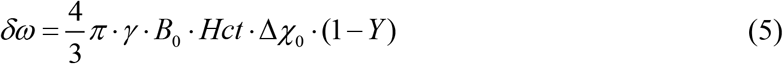

In Eq. (5), *B*_*0*_ is the MRI scanner magnetic field (3T in our data), *γ* = 2.675 · 10^8^ *rad* · *s*^−1^ · *T*^−1^ is the gyromagnetic ratio, Hct is the blood hematocrit level, Δ*χ*_0_ = 0.27 *ppm* (Spees, Yablonskiy, Oswood, & Ackerman, 2001) is the susceptibility difference between fully oxygenated and fully deoxygenated blood, and *Y* is the blood oxygenation level (Y=0 corresponds to fully deoxygenated blood and Y=1 corresponds to a fully oxygenated blood). Function *f*_*s*_ describes the signal decay due to the presence of the blood vessel network which was defined in (Yablonskiy & Haacke, 1994). Herein we use expression for the function *fs* defined in terms of hypergeometric function _1_*F*_2_ (Yablonskiy et al., 2013b) :

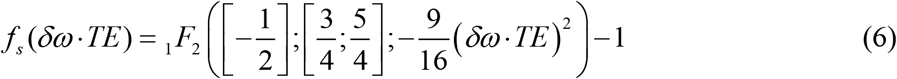

For long TE>1.5/δω this function displays linear behavior with respect to TE with a coefficient *R*2′ equal to

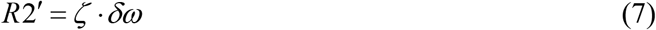

The *R*2′ is related to the GRE signal loss due to the presence of paramagnetic deoxygenated blood in veins and the pre-venous part of the capillary bed.

By making use of Eqs. (5) and (7), we can also evaluate the concentration of deoxyhemoglobin per unit tissue volume (He & Yablonskiy, 2007):

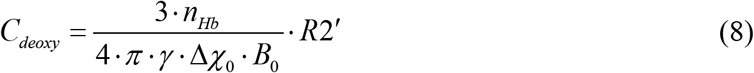

where *n*_*Hb*_ is the total intracellular Hb concentration equal to 5.5 × 10^−6^ *mol*/*mL*. Hence, parameter *k* in Eq. (2) is equal to *k* = 4 · *π* · *γ* · Δ*χ*_0_ · *B*_0_ (3 · *n*_*Hb*_).

### Image Acquisitions and biophysical model parameters estimation

A 3D multi-gradient echo sequence was used to obtain the data. Sequence parameters were: resolution 1 × 1 × 2 mm^3^ (read, phase, slab), FOV 256 mm × 192 mm, repetition time TR =50 ms, flip angle 30°, 10 gradient echoes with first gradient echo time TE_1_= 4 ms, echo spacing ΔTE = 4 ms. Additional phase stabilization echo (the navigator data) was collected for each line in *k*-space to correct for image artifacts due to the physiological fluctuations (Wen, Cross, & Yablonskiy, 2015). The total acquisition time was 11 min and 30s. After the data acquisition, the raw k-space data were read into MATLAB (TheMathWorks, Inc.) for data analysis.

Model parameters *S*_*0*_, *R*2*t*^*^, *ζ* and *δω* are obtained by fitting Eq. (3) to experimental data on a voxel-by-voxel basis using non-linear least square (NLLS) algorithm. The function Fmacro(TE) is precomputed using VSF method (Yablonskiy et al., 2013b) with fast algorithm (Wen et al., 2020).

To mitigate the adverse effects of the noise on parameters estimation we adopted a two-step fitting procedure (D. A. Yablonskiy, J. Wen, S. Kothapalli, & A. L. Sukstanskii, 2021). At the first step, we evaluate all parameters using NLLS algorithm. At the second step, we calculate mean δω for each ROI defined using FreeSurfer segmentation and then apply NLLS with fixed δω to estimate *S*_*0*_, *R*2*t*^*^ and *R*2′ = *ζ* · *δω*. To further reduce noise influence on the data, we apply the Hanning filter in the image domain to the *R*2*t*^*^ and *R*2′ maps to use as reference data for training our model.

### DANSE Architecture and Training

In this study we use deep convolutional neural network model based on 2D U-Net architecture (similar to (Torop et al., 2020)) as illustrated in Fig. 1. The U-Net architecture (Çiçek, Abdulkadir, Lienkamp, Brox, & Ronneberger, 2016; Ronneberger, Fischer, & Brox, 2015) consists of contracting path and expanding path with two paths being symmetrical. Each contracting path includes a series of convolutions, each followed by a rectified linear unit (ReLU) and a max pooling operation to encode the local information while the expanding path uses sequence of up-convolutions with high-resolution feature concatenations. The high resolution features from the contracting path were concatenated with the outputs of expanding path for future reuse and compensating resolution loss in the final output. The neural network model was implemented in TensorFlow (Abadi et al., 2016). The network weights were learned by minimizing the *L*_*1*_ loss between the prediction and the ground truth reference using Adam (Kingma and Ba, 2014) optimizer with an initial learning rate of 8e-4 and a batch size of 4. DANSE training was done by splitting the aforementioned 26 data into 18 datasets (≈70%) for training, 4 for validation (≈15%) and 4 for testing (≈15%). The model was trained from scratch with random weights initialization. Such multi-scale structure leads to a large receptive field of the CNN that is effective for removing globally spread imaging artifacts typical in MRI (Han et al., 2018; Krizhevsky, Sutskever, & Hinton, 2012).

**Figure 1:**
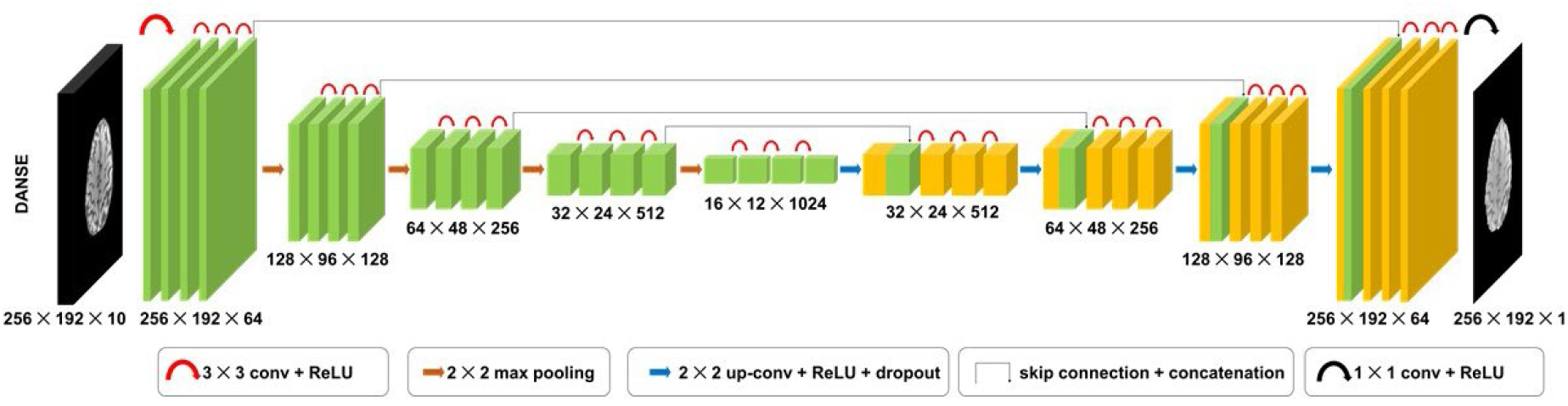
A fully supervised Deep-Learning-Based Accelerated and Noise-Suppressed Estimation (DANSE) method for generating qGRE metrics associated with human brain neuronal structure and hemodynamic properties. Figure represents a schematic structure of the convolutional neural network model based on 2D U-Net architecture with four-level depth. It takes 10 qGRE magnitude images (corresponding to 10 gradient echoes) to reconstruct a single map representing one of the qGRE parameters (S_0_, *R*2*t*^*^ and *R*2′).

In this study, we have trained the DANSE model separately for each map of the three biophysical model parameters *p*_*n*_, i.e. *S*_*0*_, *R*2*t*^*^ and *R*2′. The ground truth maps for each parameter were generated by fitting the biophysical model in Eq. (3) as discussed in previous section. Our model processes data from individual spatial slices extracted from 3D qGRE MRI data. The 3D image of the whole brain is then obtained by concatenating the outputs of the DANSE applied slice-by-slice. We represent the magnitudes of the measured 2D qGRE images of *M* gradient echo times as follows:

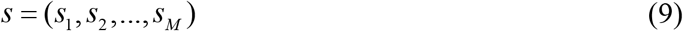

where each vector *s*_*m*_ in *s* (*m* = 1, 2, …, *M*) denotes a vectorized 2D image representing the magnitude of the data for one of the gradient echo times.

Figure 2 demonstrates a pictorial schematic design for the prediction of brain maps corresponding to each true map. The ground-truth of the desired brain maps (*p*_*n*_) are represented as: *p*_1_ = *S*_0_, *p*_2_ = *R*2*t*^*^ and *p*_3_ = *R*2′. Let 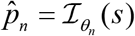 denote our deep CNN architecture which estimates the values 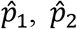 and 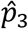 of corresponding true values of *p*_*n*_ determined by the biophysical model from the qGRE signal *s*. The vector *θ*_*n*_ denotes the trainable set of weights for parameter *n* in the CNN. As illustrated in Fig. 1 DANSE takes 10-echo magnitude data and produces high quality maps of biophysical parameters free from field inhomogeneity artifacts. It is to be noted that, we train DANSE for each brain parameter map separately and for each case we get 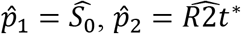, and 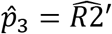 as the estimated parameters from DANSE model by minimizing the empirical loss: 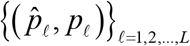, over the training set consisting of *L* 2D-slices. Common choices for 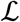 include the Euclidean and the ℓ_1_ distances. The equation can be expressed as follows:

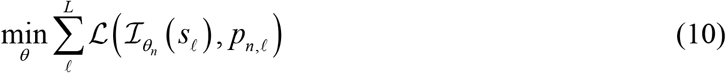

where *p*_*n,l*_ represents maps of one of the reference biophysical parameters {*p*_1_, *p*_2_, *p*_3_} for each slice *l*. To solve the minimization problem the stochastic gradient-based optimization algorithms such as Adam (Bottou, 2012; Kingma & Ba, 2014) is used. Once the optimal set of parameters 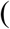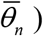 has been learned on the training data, the operator 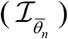 is applied to solve the computational problem on previously unseen data. It is important to note that the function *Fmacro* is required only for generating ground truth maps of biophysical parameters but it is not used either for model training or testing. Each 3D qGRE MR dataset is normalized to the mean brain signal of first gradient echo of corresponding dataset before feeding the slices to the DANSE model. This step is intended to make our DANSE model compatible with MR data obtained from different scanners.

**Figure 2:**
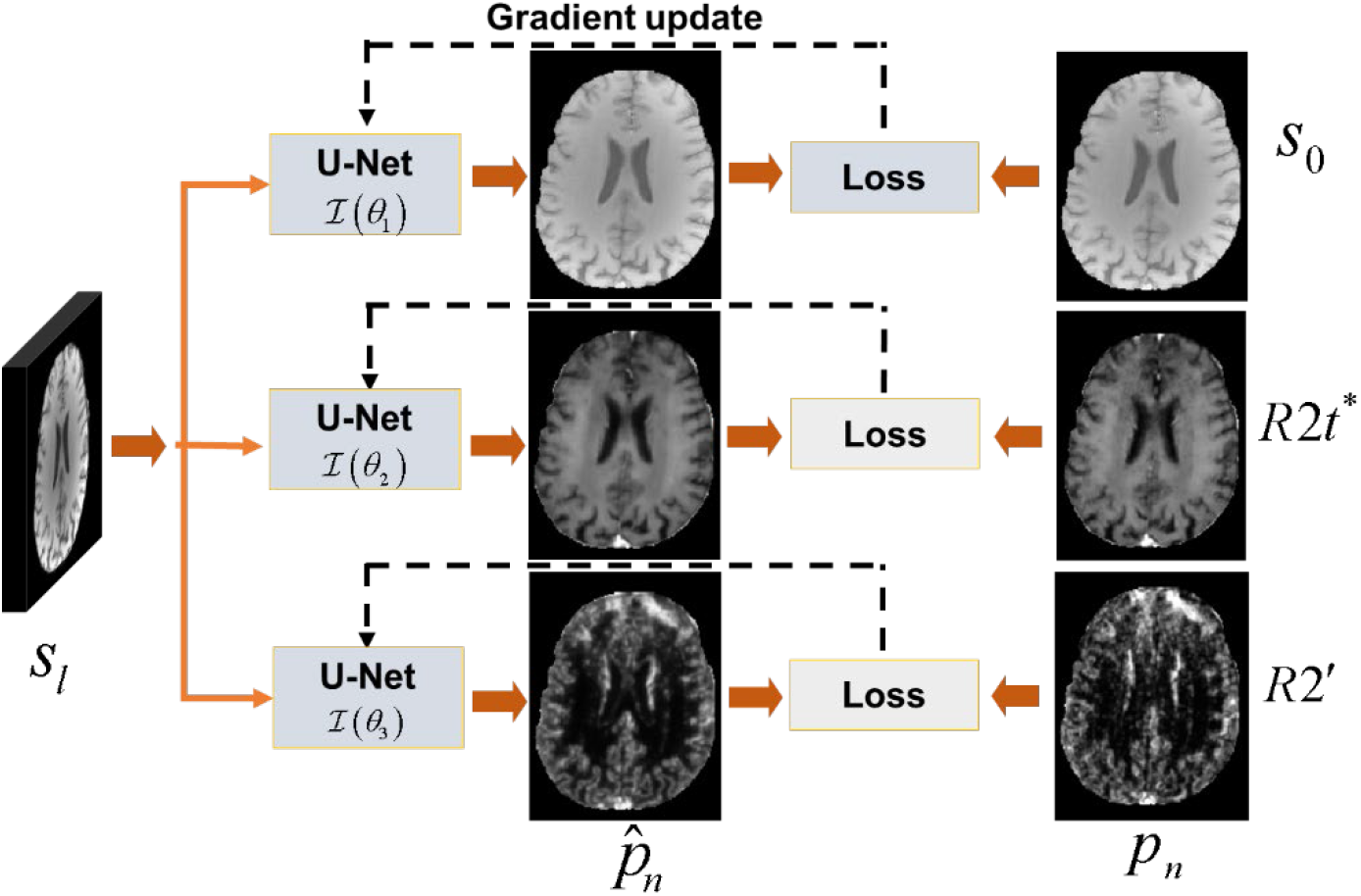
DANSE supervised training model. For each map of the parameters *p*_*n*_, the individual model’s weights *θ*_*n*_ are optimized for the loss, so that 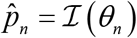 is close to the corresponding ground-truth maps of *p*_*n*_.

### Noise Sensitivity of DANSE model

To address the question of how noise affects our model, we have generated simulated data by using maps of parameters *S*_*0*_, *R*2*t*^*^, *ζ* and *δω* and values of F_macro_(TE) obtained from 26 subjects. For the ground truth we used 26 simulated image data-sets obtained by substituting values of these parameters in Eq. (3) thus obtaining 26 new data-sets, each containing 10 3D images corresponding to 10 TEs. The noisy data were generated by adding noise corresponding to SNRs 50, 100, 175 and 200 to the simulated data. The random noise values were drawn from the standard normal distribution generated by a built-in function in MATLAB.

The performance of DANSE model is validated by training on noisy simulated data with different noise levels. More particularly, the training dataset consists of datasets with varying SNR levels. We made four sets where each having 7 datasets with SNRs 50, 100 and 175 while remaining 5 were with SNR 200. This was done to make our model more robust to different noise levels. Then we tested our model on data with SNRs 50, 75, 125, 175 and 200.

To quantitatively assess the performance of DANSE model, we use *relative error* (*RE*) and *structural similarity index (SSIM)* between the estimated result (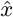) with its reference *x* values. SSIM is a perceptual metric which assesses the perceived quality of an image based on luminance, contrast and structure. It can be computed using its definition in (Z. Wang, Bovik, Sheikh, & Simoncelli, 2004). *RE* is defined as:

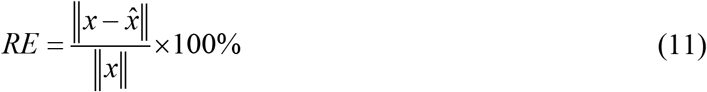

where 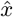 and *x* are the estimated result from DANSE and its corresponding ground truth reference, respectively. Here, **‖**·**‖** denotes the standard Euclidean norm.

## Results

Our DANSE model generates the qGRE parameters’ maps free from magnetic field inhomogeneity artifacts directly from the GRE MRI magnitude data reflecting qGRE signal decay. It is important to note that the reference images were generated by considering precomputed magnetic field inhomogeneities using voxel spread function (Yablonskiy et al., 2013a) but model application does not require any information about field inhomogeneities. Two examples of GRE magnitude images are shown in Figure 3 which exhibit reduction in the GRE image intensity due to the R2* signal decay with increase in gradient echo time TE. Figure 4 illustrates two examples of estimated brain maps for each parameter by DANSE model and their corresponding ground truths. It can be clearly noticed that our DANSE model estimates high quality, brain maps as compared to the reference brain maps while preserving the contrast for all maps.

**Figure 3.**
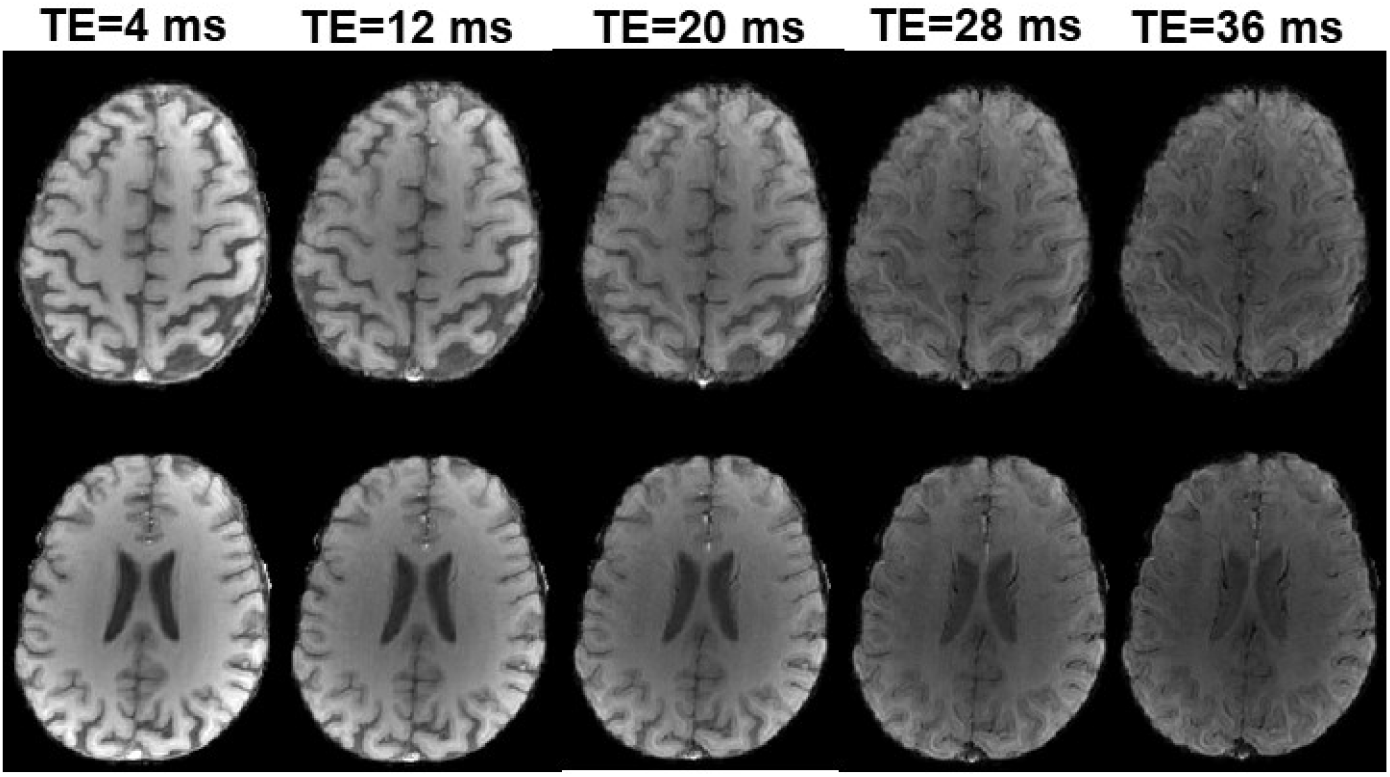
Two examples of *in-vivo* magnitude images exhibiting signal decay with gradient echo time TE increasing from 4 ms to 36 ms. Only every other gradient echo image is shown while actual 10 gradient echo images are collected with 4-ms intervals.

**Figure 4.**
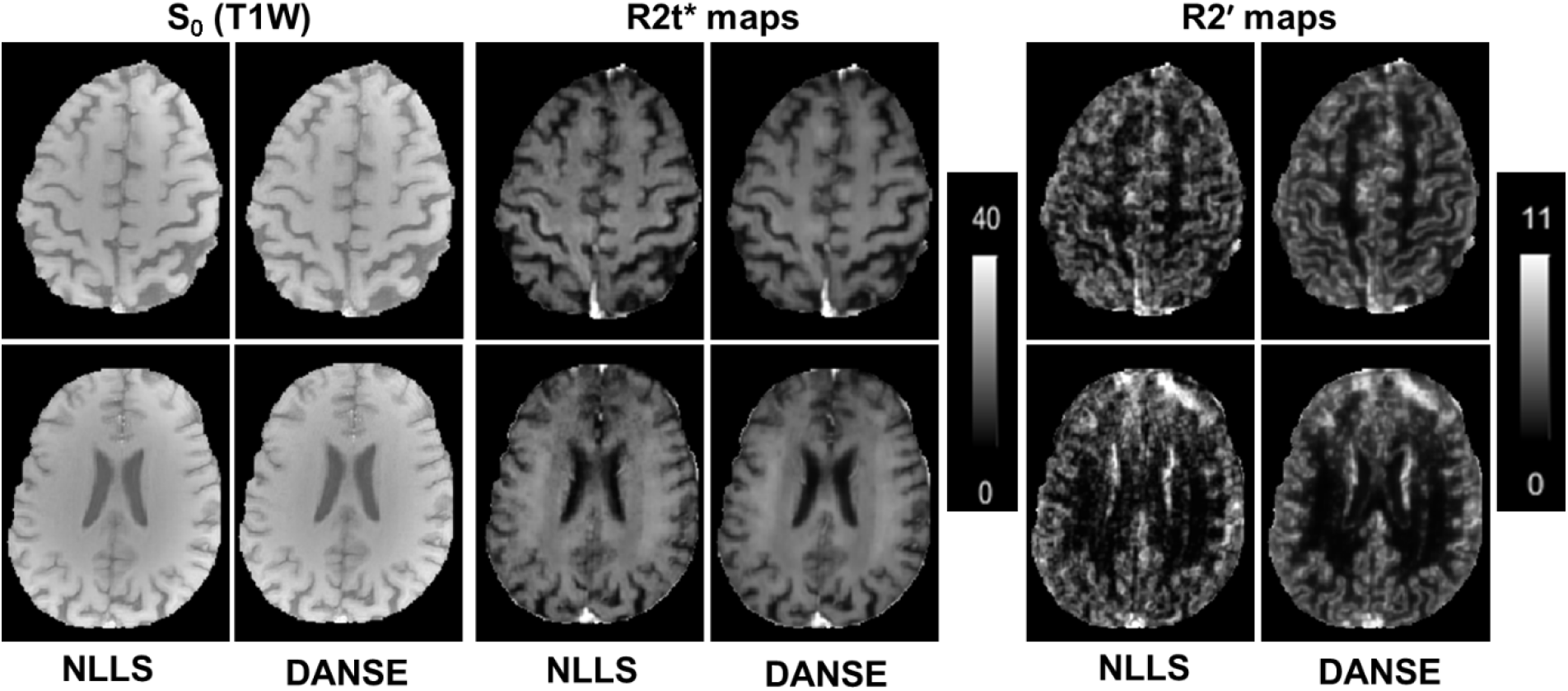
Two examples of qGRE parameter maps estimated by DANSE model and NLLS fitting from in vivo data. Columns 1, 3 and 5 show *S*_0_, *R*2*t*^*^ and *R*2′ from NLLS approach, respectively while columns 2, 4 and 6 show the corresponding maps from DANSE method. Relative Differences between DANSE and NLLS results are 2%, 3% and 11% for *S*_0_, *R*2*t*^*^ and *R*2′, respectively.

Figure 4 shows two examples of *S*_0_, *R*2*t*^*^ and *R*2′ maps obtained by DANSE and NLLS methods from *in-*vivo data. Relative differences between DANSE and NLLS results are 2%, 3% and 11% for *S*_0_, *R*2*t*^*^ and *R*2′, respectively. While images look quite similar, the DANSE results appear less noisy. Moreover, we have calculated SSIM in order to assess the image quality where the first row values for *S*_0_, *R*2*t*^*^ and *R*2′ (columns 2, 4 and 6) are 0.99, 0.97 and 0.93, respectively while the second row SSIM values for the same maps are 0.99, 0.94 and 0.90, respectively.

To quantitatively assess the noise sensitivity of DANSE model we have trained it by generating synthetic data based on biophysical model and adding noise with different level corresponding to SNRs 50, 100, 175 and 200. Figures 5 and 7 present examples of *R*2*t*^*^ and R2′ maps obtained by means of NLLS and DANSE method for two slices from simulated data with different noise levels (SNR 50, 125 and 200). Figures 6 and 8 show the plots comparing errors in *R*2*t*^*^ and R2′ parameters estimations by means DANSE and NLLS approaches. It can be observed that the DANSE model significantly outperforms the NLLS method for all noise variations.

**Figure 5.**
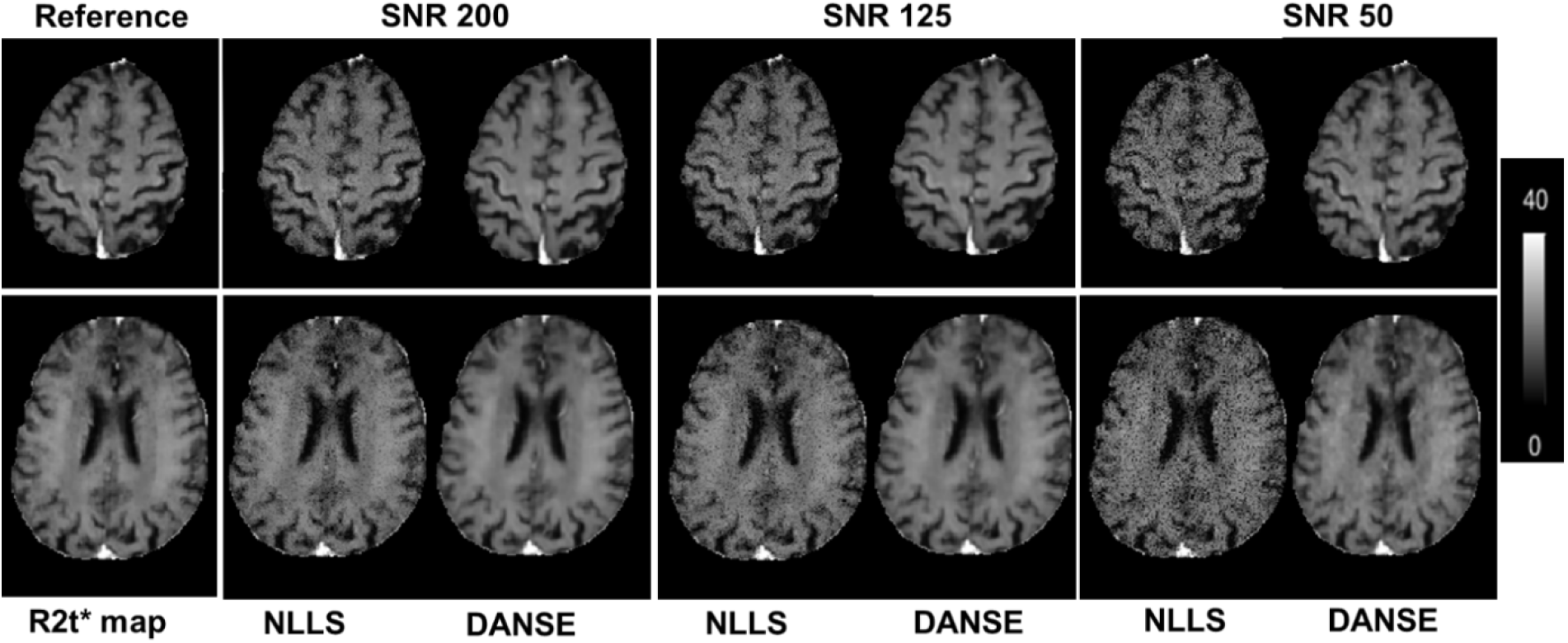
Two examples of R2t* maps estimated by NLLS and DANSE model from simulated data with different noise levels. Reference (noiseless) R2t* maps are also shown in the left column. While effect of noise on NLLS results is clearly visible for all SNRs and is increasing with SNR reduction, the DANSE model produces results with a small error (about 3%) practically similar to the reference R2t* maps independent of the noise level. This result is quantitatively illustrated in Fig. 6.

**Figure 6.**
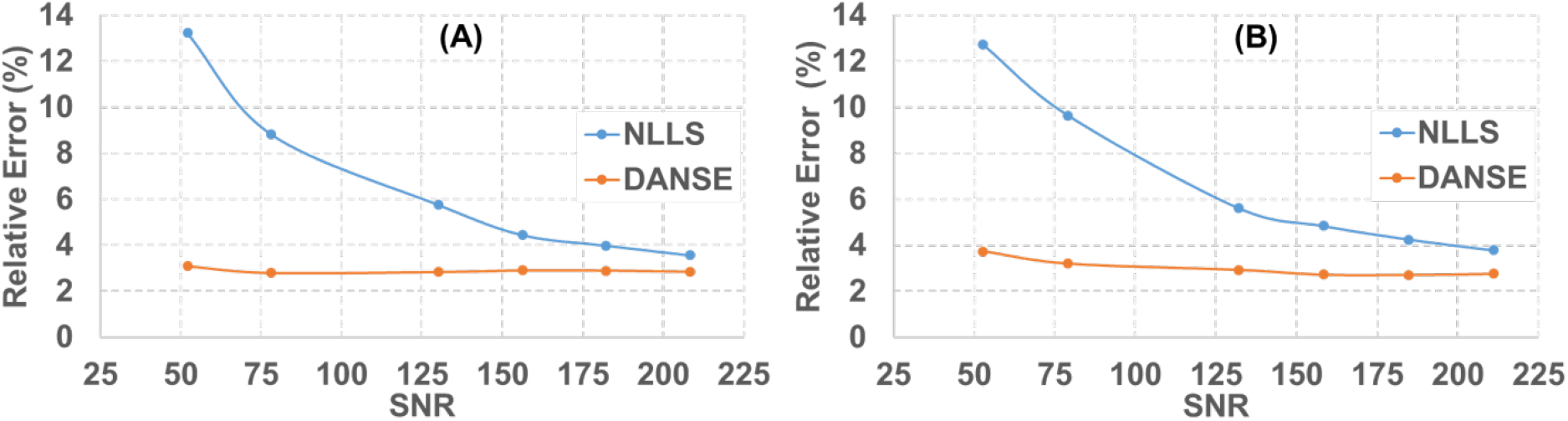
Plots represent percentage of errors between estimated *R*2*t*^*^ by DANSE model and NLLS results as function of SNR for two slices presented in Fig. 5. (A) corresponds to the upper row in Fig. 5 and (B) to the lower row.

**Fig. 7.**
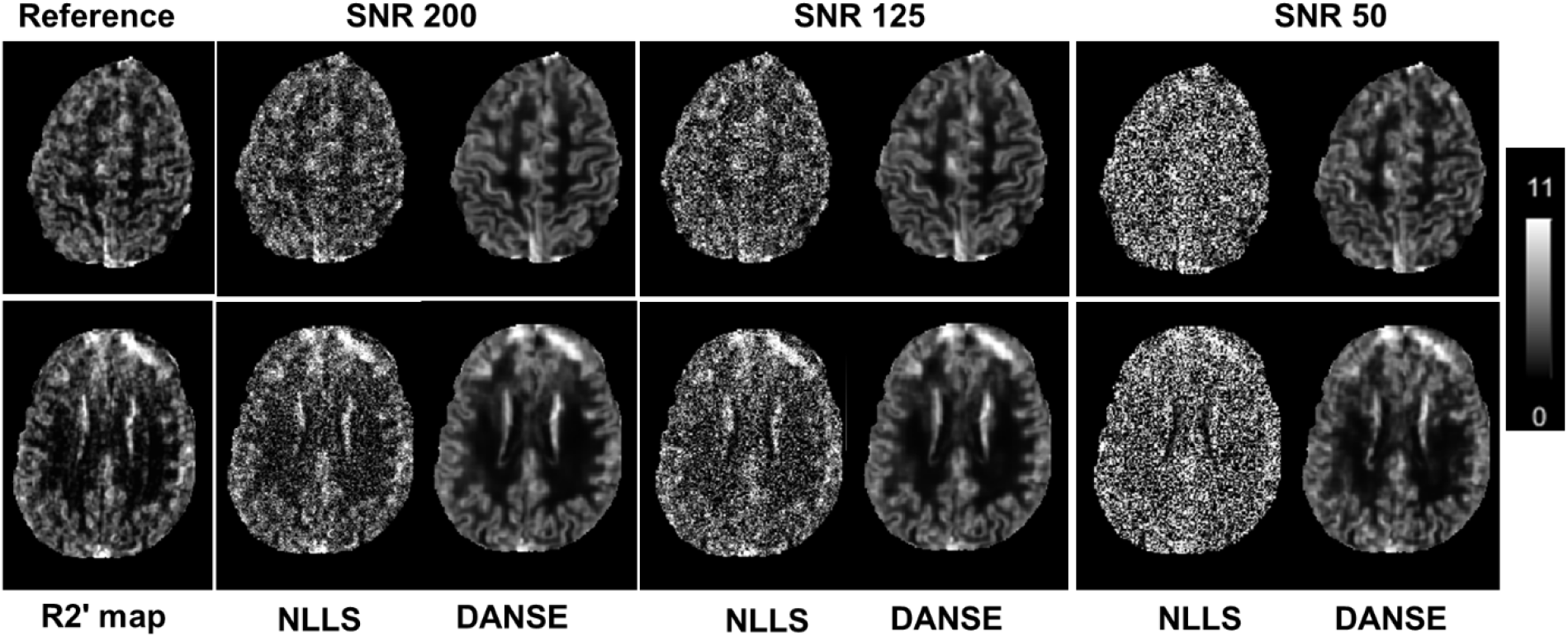
Examples of *R*2′ maps from NLLS and DANSE methods for two slices from simulated data with different noise levels (SNR 50, 125 and 200). Reference (noiseless) R2′ maps are also shown in the left column. While effect of the noise on NLLS results is significant even for SNR 200 and becomes overwhelming for lower SNR, the DANSE model produces practically independent of SNR level results with the error less than 20%. These data are quantitatively illustrated in Fig. 8.

**Fig. 8.**
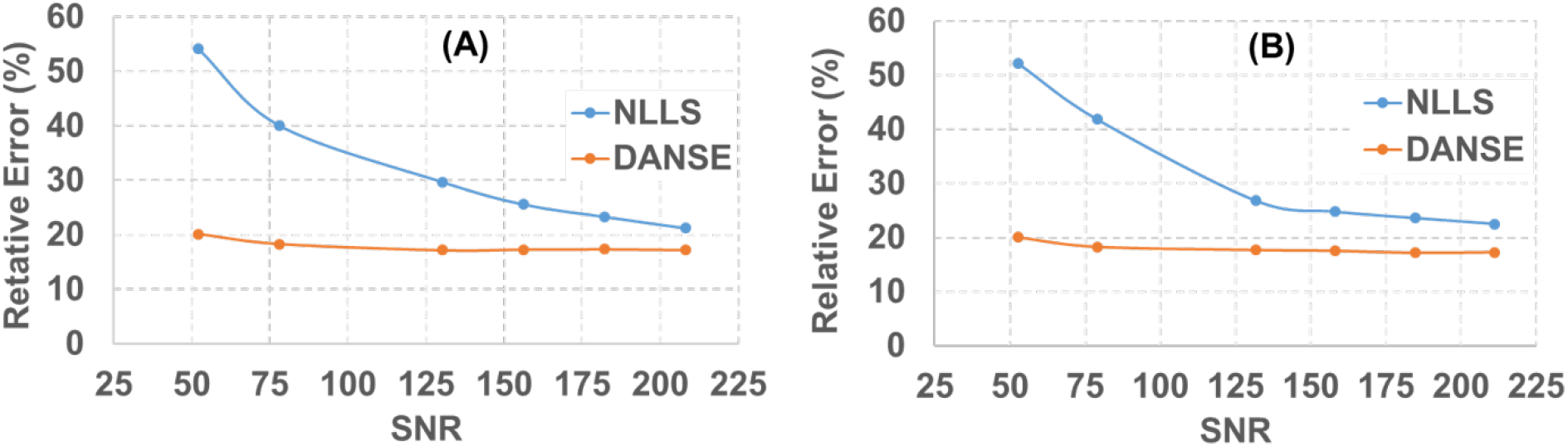
Plots represent percentage of errors between estimated *R*2′ by DANSE model and NLLS results as function of SNR for two slices presented in Fig. 7. (A) corresponds to the upper row in Fig. 7 and (B) to the lower row.

## Discussion

Deep learning is becoming a mainstream approach for analyzing medical images. Combining deep learning with biophysical model relating MRI data to biological tissue microstructure (Torop et al., 2020) allows fast generating of quantitative images with enhanced tissue-specific information than so-called weighted images mostly used in clinical practice.

In this paper we present a Deep-Learning-Based Accelerated and Noise-Suppressed Estimation (DANSE) method that permits calculation of quantitative Gradient Recalled Echo (qGRE) MRI metrics associated with human brain neuronal structure and hemodynamic properties in a matter of seconds as compared with hours required by using non-linear least square algorithms. DANSE method uses a supervised convolutional neural network (CNN) approach for fast and robust estimation of high-quality T1-weighted images (*S*_0_), as well as *R*2*t*^*^ and *R*2′ maps generated from qGRE MRI data.

The *R*2*t*^*^ metric of qGRE signal describes the part of the signal decay resulting from water molecules (major source of MRI signal) interaction with cellular and extracellular components of biological tissues and is mostly associated with the cortical brain tissue neuronal density (Wen et al., 2018). It is sensitive to pre-atrophic neurodegeneration even in preclinical stages of Alzheimer disease (Kothapalli et al., 2021; Zhao et al., 2017). It has also proved useful in applications to multiple sclerosis (Xiang et al., 2019) and traumatic brain injury (Astafiev et al., 2019). The *R*2*t*^*^ metric can also provide quantitative information on non-heme iron in basal ganglia (D. A. Yablonskiy, J. Wen, S. V. V. N. Kothapalli, & A. L. Sukstanskii, 2021).

The *R*2′ metric of qGRE signal reflects the MRI signal decay due to the BOLD (Blood Oxygenation Level Dependent) effect (Ogawa et al., 1992) and provides important information on tissue metabolic properties which are important in understanding normal brain operation and pathophysiology of different neurological disorders such as, Alzheimer’s disease (Iadecola, 2004, 2005; Vlassenko et al., 2010), Parkinson’s disease (Beal, 1998; Schapira, 1998), stroke (Derdeyn et al., 2002), psychiatric diseases (Mamah et al., 2015) etc.

In the paper, we have built DANSE model that can act on in vivo data and then also tested model resilience to noise. For in vivo applications, the reference *S*_0_ images, *R*2*t*^*^ and *R*2′ maps for training the model were generated by means of bio-physical model, Eq. (3), that unveils MRI signal properties related to biological tissue microstructure. Figure 4 shows two examples of *S*_0_ images, *R*2*t*^*^ and *R*2′ maps obtained from in vivo data with DANSE model and by means of voxel-by-voxel NLLS fitting. DANSE approach generates images reproducing all the features present in the NLLS-generated images but with reduced effects of noise in the data. In this context, it is to be noted that DANSE method produces the brain maps (*S*_0_, *R*2*t*^*^ and *R*2′) significantly faster than the NLLS fitting procedure. DANSE takes 30 seconds to compute the parameters for the entire brain while NLLS takes several hours to estimate the same parameters on modern computers. It is important to note that DANSE model recognizes the contributions of magnetic field inhomogeneities simply from the intensity inhomogeneity in the qGRE magnitude images and produces high quality brain maps similar to NLLS method free of magnetic field inhomogeneity artifacts. At the same time, the NLLS procedure estimates the contribution of field inhomogeneity effects by additional calculation of the *F*_*macro*_ in Eq. (3) by means of the voxel-spread-function approach (Yablonskiy et al., 2013a).

To assess the noise resilience of DANSE model we have trained it by generating synthetic data based on biophysical model and adding noise with different level corresponding to SNR of 50, 100, 175 and 200. Figures 5 and 7 show the comparison between qualitative outputs from two slices for maps of *R*2*t*^*^ and *R*2′, respectively from DANSE and NLLS approaches with different levels of SNR. It is quite evident that DANSE model works well even in environments with high noise levels and produces clean images with very low noise contamination as compared with NLLS method. While the DANSE model produces practically independent of SNR level results with the error less than 20% for *R*2′ and about 3% for *R*2*t*^*^, the effect of the noise on NLLS results is significant even for SNR 200 and becomes overwhelming for lower SNR. Notably, for SNR 50, the errors in estimating *R*2*t*^*^ and *R*2′ by DANSE method are three times smaller than errors of NLLS approach.

## Conclusion

The DANSE method proposed in this paper allows fast reconstruction of quantitative qGRE metrics related to biological tissue cellular structure (*R*2*t*^*^) and hemodynamic properties (*R*2′), along with T1-weighted images (S_0_). The key features of DANSE model include:

1. The DANSE model requires only magnitude images while producing *S*_0_, *R*2*t*^*^ and *R*2′ maps *free of magnetic field inhomogeneity artifacts*. This was possible because the DANSE model was trained using biophysical model that accounted for the adverse effects of magnetic field inhomogeneities (the factor *F*_*macro*_*(TE)* in Eq. (3)) by means of the voxel spread function (VSF) method (Yablonskiy et al., 2013b).
2. The DANSE model has significant “*noise-suppressing*” advantage as compared with the standard NLLS reconstruction that is especially evident for data with low SNR. For example, for SNR 50, the errors in estimating *R*2*t*^*^ and *R*2′ by DANSE method are three times smaller than the errors of NLLS approach.
3. The DANSE model offers significant advantage in the speed of image reconstruction – it takes only 30 seconds to compute the qGRE metrics (*S*_0_, *R*2*t*^*^ and *R*2′) for the entire brain with DANSE while NLLS takes several hours to estimate the same parameters on modern computers.

Since the qGRE technique has already proved useful in studying healthy brain aging (Zhao et al., 2016), psychiatric diseases (Mamah et al., 2015), loss of neurons at early stages of Alzheimer disease (Kothapalli et al., 2021; Zhao et al., 2017), multiple sclerosis (Xiang et al., 2019) and traumatic brain injury (Astafiev et al., 2019), the DANSE method, by providing fast and robust approach for generating qGRE metrics associated with human brain neuronal structure and hemodynamic properties, may open doors to broad qGRE applications in clinical practice.

## ACKNOWLEDGMENTS

This work was supported by NIH/NIA grant R01AG054513, NSF CAREER award CCF-2043134, and Marilyn Hilton Award for Innovation in MS Research.

